# Comprehensive molecular features of polycystic ovary syndrome revealed by transcriptome analysis of oocytes and cumulus cells

**DOI:** 10.1101/2021.01.30.428778

**Authors:** Jie Li, Haixia Chen, Mo Gou, Chenglei Tian, Huasong Wang, Xueru Song, David. L. Keefe, Xiaohong Bai, Lin Liu

**Author notes:** Equal contribution (Jie Li, Haixia Chen).

## Abstract

PCOS is typically characterized by polycystic ovarian morphology, hyperandrogenism, ovulatory dysfunction and infertility. Furthermore, PCOS patients undergoing ovarian stimulation have more oocytes, however, poor quality of oocytes lead to lower fertilization and implantation rates, decreased pregnancy and increased miscarriage rates. Our study suggests that global gene expression and cell to cell interactions of oocytes and CCs are significantly altered in women with PCOS. Noticeably, genes related to microtubules such as TUBB8 and TUBA1C are abnormally highly expressed in PCOS oocytes, reducing oocyte quality. The pattern of transposable element expression distinguishes PCOS from Control oocytes, implying the role of transposable elements in the occurrence of PCOS.

## Introduction

Polycystic ovary syndrome (PCOS) is the most common endocrinopathy in women of reproductive age, with a prevalence of about 10% ^1^. The syndrome is typically characterized by polycystic ovarian morphology, hyperandrogenism and ovulatory dysfunction. Additional clinical features include insulin resistance, obesity, type 2 diabetes (T2D) and infertility ^2^. Furthermore, a recent study shows that daughters of women with PCOS are more often diagnosed with PCOS. PCOS phenotype also can be transgenerationally transmitted across offspring of female mice ^3^. Previous GWAS analysis identified 11 loci associated with PCOS and candidate genes at these loci, which were related to clinical manifestations of PCOS, such as infertility, insulin resistance, T2D and others ^4,5^ These studies facilitated understanding of the etiologic factors accounting for PCOS. Microarray or RNA-sequencing analysis of oocytes and/or granulosa cells or cumulus cells in women with PCOS has provided insights into understanding of PCOS ^6–10^.

Notably, at least 40% of the human genome derives from transposable elements. Yet, transcription of TEs in PCOS patients remains to be comprehensively investigated. TEs are categorized into two features-DNA transposons accounting for 3% of TEs and retrotransposons represents 90% of TEs ^11^, and retrotransposons are further divided into five orders including LTR, DIRS, PLE, LINE and SINE ^12^. Endogenous retroviruses (ERVs), as a superfamily of LTR retrotransposons, can copy and paste their own DNA into the genome ^13,14^ TEs also play essential roles in transcriptional modulation, and specific ERV families are transcribed during human preimplantation development, which is stage specific ^15^. Somatic retrotransposons can alter the expression of protein-coding genes differentially expressed in the human brain ^16^. Malki et al. proposed that fetal oocyte attrition (FOA) selects oocytes with low activity of a specific retrotransposon, Long Interspersed Nuclear Element 1 (L1). L1 activity, therefore, may be involved in controlling the size and quality of mammalian ovarian oocytes reserve ^17^ Yet, L1 methylation levels are only slightly changed in cumulus cells of oocytes from patients with PCOS ^18^. Dysregulated TEs are involved in gene mutation and the occurrence of a number of human diseases, including malignancies, neurological disease and normal aging ^19^.

In this study, we systematically analyzed TE and global gene expression in oocytes and CCs from PCOS patients and compared them with Controls. We identified new candidate genes and TEs underlying PCOS, which may serve as biomarkers of PCOS.

## Results

### Global gene expression profile of oocytes distinguishes PCOS from Control

The study included five PCOS patients and five Controls, and clinical characteristics were shown in Supplementary Table SI. PCOS patients exhibited a significantly higher LH levels (7.71 ± 1.11) than did Controls (3.93 ± 0.65). Compared to Controls, PCOS patients had lower FSH levels (4.91 ± 0.46), elevated testosterone (52.87± 3.93) and E2 levels (5813.53 ± 298.03). In addition, the number of antral follicles and oocytes retrieved from PCOS patients (31.40 ± 1.69 and 34 ± 7.62, respectively) were significantly more than that of Controls (16.80 ± 3.40 and 15.20 ± 2.78). Next, we performed RNA sequencing of six GV oocytes and four CC samples from the same PCOS patients or Controls. The sequencing depth of libraries for all oocyte was sufficient to ensure the accuracy and consistency of subsequent analysis. After quality control and filtration of the RNA-seq data, we identified 24,251 genes expressed in oocytes and 42,725 genes in CC samples. The obtained RNA-seq normalized data were used for dimensional reduction analysis (t-SNE) ^20^ by an unsupervised approach, showing that there was a clear separation between oocytes and matched CCs with or without PCOS (Supplementary Fig. S1F). Interestingly, we found distinct PCOS-specific clustering of oocytes, whereas clustering of CCs displayed some overlap. Meanwhile, PerMANOVA analysis further confirmed significant differences in global gene expression of oocytes or CCs between PCOS and Control group (Supplementary Table SIV, *P* =0.004, *P=* 0.03). Hence, oocytes and CCs had different gene expression patterns in the occurrence of PCOS.

To explore the pattern of gene expression in oocytes from PCOS, we applied t-SNE to normalize expression data by using unsupervised clustering. PCOS and Control clusters could be noticeably differentiated (Fig. 1A). Next, we examined differentially expressed genes (DEGs) between PCOS (1,433 genes upregulated and 1,322 genes downregulated) and Control oocytes (Fig. 1B; Supplementary TableV). Although these data were only generated from three PCOS patients and three Controls, such consistent gene expression patterns among them shown in the heatmap demonstrated robustness of our RNA-seq analysis as well as minimal variations among patients. By GO (Gene Ontology) analysis, crucial functions *(P* < 0.05) enriched by upregulated DEGs are displayed (Supplementary Fig. S1G). It is noteworthy that genes associated with the function of chromatin, microtubule, cytoskeleton, and actin were upregulated in oocytes from PCOS women (Fig. 1C, Fig.1D). Among genes involved in microtubule-based process, *TUBB8* and *TUBA1C* exhibited the highest expression levels in PCOS oocytes. Immunofluorescence (IF) of oocytes confirmed TUBB8 and TUBA1C overexpression in oocytes from women with PCOS (Fig. 1E). It has been found that TUBB8 plays a key role in meiotic spindle assembly and maturation in human oocytes and mutations in TUBB8 leads to oocyte maturation arrest ^21,22^.

**Figure 1.**
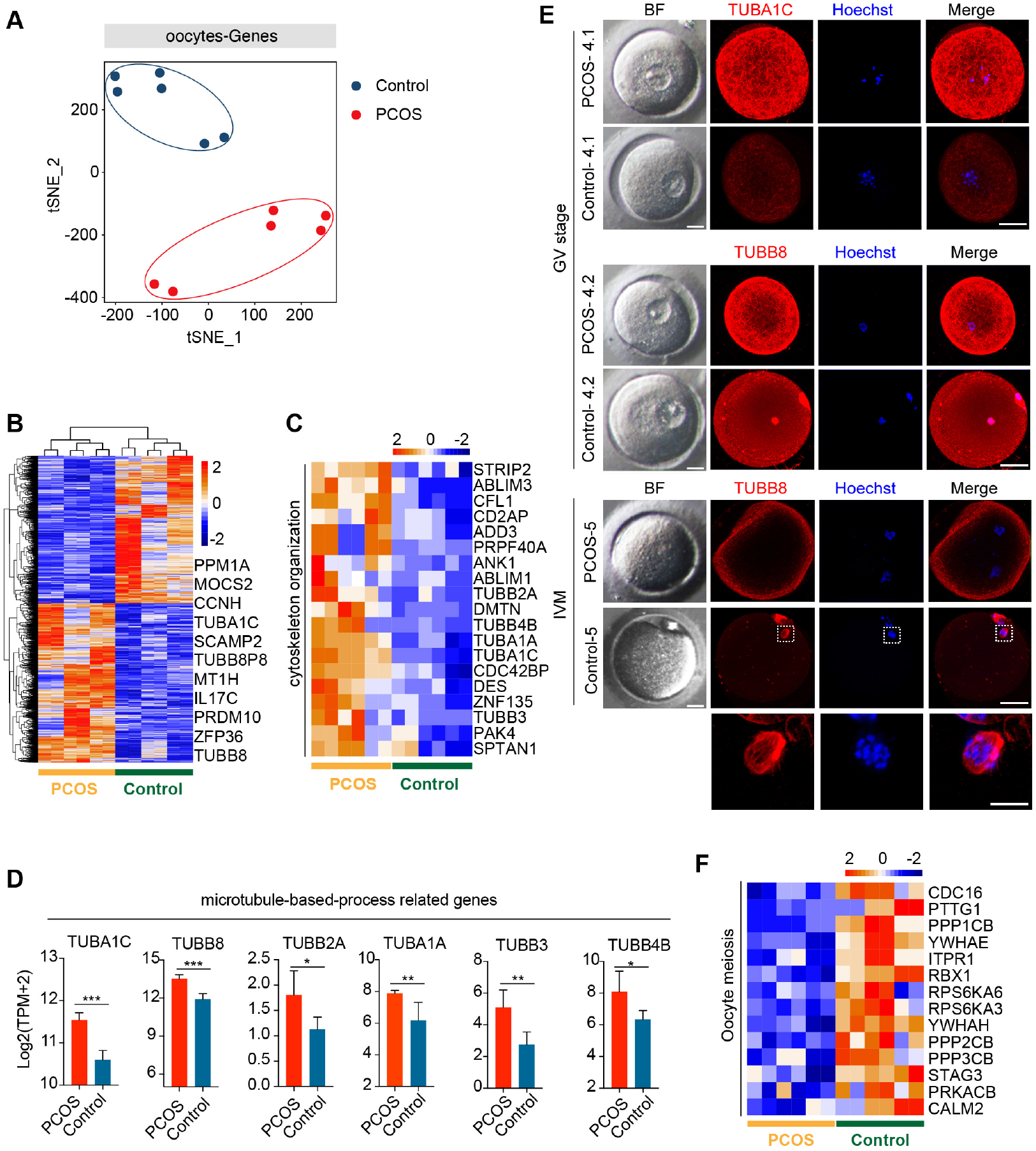
Global gene expression profile distinguishes PCOS from Control oocytes. (**A**) Visualization of the gene expression of six oocytes by t-SNE, which are clustered into two subpopulations including Control group and PCOS group, red and blue points represent PCOS and Control oocytes, respectively. (**B**) Heatmap displaying differentially expressed genes (DEGs) in oocytes between PCOS and Control. The number of up-DEGs is 1,433 and down-DEGs is 1,322. The color key from blue to red indicates the relative gene expression levels from low to high, respectively. (**C**) Heatmap of genes involved in cytoskeleton organization that were upregulated in PCOS oocytes. (**D**) Six genes associated with microtubule based process. Red and blue bar represent PCOS and Control, respectively. Gene expression levels are represented by log_2_ [TPM+2], and data represents mean ± SD. n = 3 (participants). *p < 0.05, **p < 0.01, ***p < 0.001, ns, not significant. (**E**) Bright-field and immunofluorescence images of TUBA1C and TUBB8 in normal and PCOS oocytes at the GV stage or MII stage after IVM. Upper panel, oocytes (PCOS and Control) at GV stage and two PCOS oocytes or two Control oocytes were obtained from the same person. Lower panel, oocytes (PCOS and Control) after in-vitro maturation. Hoechst (blue) was used to counterstain and visualize DNA. Scale bar represents 20 μm. Magnifications of the spindle regions are shown at the bottom and the scale bar is 10 μm (bottom). (**F**) Heatmap of genes associated with oocyte meiosis downregulated in PCOS oocytes.

Meanwhile, the pivotal function (*P* < 0.05) enriched by downregulated DEGs was also revealed (Supplementary Fig. S1H). By KEGG (Kyoto Encyclopedia of Genes and Genomes) analysis, significantly downregulated signaling pathways in PCOS oocytes included MAPK, mTOR and FOXO signaling pathways (Supplementary Fig. S1I). MAPK signaling pathway is important for the cell cycle of human oocytes ^23,24^, and mTOR–eIF4F pathway spatiotemporally regulates the translation of mammalian oocytes in meiosis ^25^. Upregulated genes also were enriched for signaling pathways (P < 0.05) of spliceosome and gap junctions (Supplementary Fig. S1J). Gap junctions play an important role in communication between oocytes and CCs, and breakdown of gap junction in the ovarian follicle induces recommencement of meiosis ^26,27^.

Based on the above GO and KEGG analysis of oocytes from PCOS, we considered that disordered expression of several signaling pathways and dysfunction of cytoskeleton, specifically microtubules, may result in meiotic abnormality in oocytes from patients with PCOS. Indeed, while control oocytes reached MII with clearly visible, barrel-shaped spindle with well-aligned chromosomes, PCOS oocytes displayed maturation arrest and no spindles, as well as disrupted TUBB8 (Fig. 1E). Genes involved in meiosis also were downregulated in PCOS oocytes (Fig. 1F). Taken together, our data revealed that the aberrant gene expression profile, including cytoskeleton, microtubule and meiosis, likely impairs the developmental competence of oocytes from patients with PCOS.

### Gene expression pattern of CCs during the occurrence of PCOS

Intimate communication between oocytes and CCs plays a crucial role in follicle development and oogenesis ^28^. We also performed RNA-seq analysis of CCs from the same patients who donated the oocytes. All CCs were clustered into two groups, depending on whether suffering from PCOS or not (Fig. 2A). The clear separation of t-SNE pattern indicated a pronounced difference in the transcriptome of PCOS and Control CCs. We then characterized differentially expressed genes (DEGs) between PCOS (704 upregulated genes and 1,091 downregulated genes) and Control CCs (Fig. 2B; Supplementary Table VI). GO terms (biological process) enriched by upregulated DEGs were found in PCOS CCs (Supplementary Fig. S2A). Several genes associated with positive regulation of apoptotic process demonstrated an increased expression levels in PCOS CCs (Fig. 2C). Also, genes associated with positive regulation of GTPase activity were overexpressed in CCs from women with PCOS (Fig. 2D).

**Figure 2.**
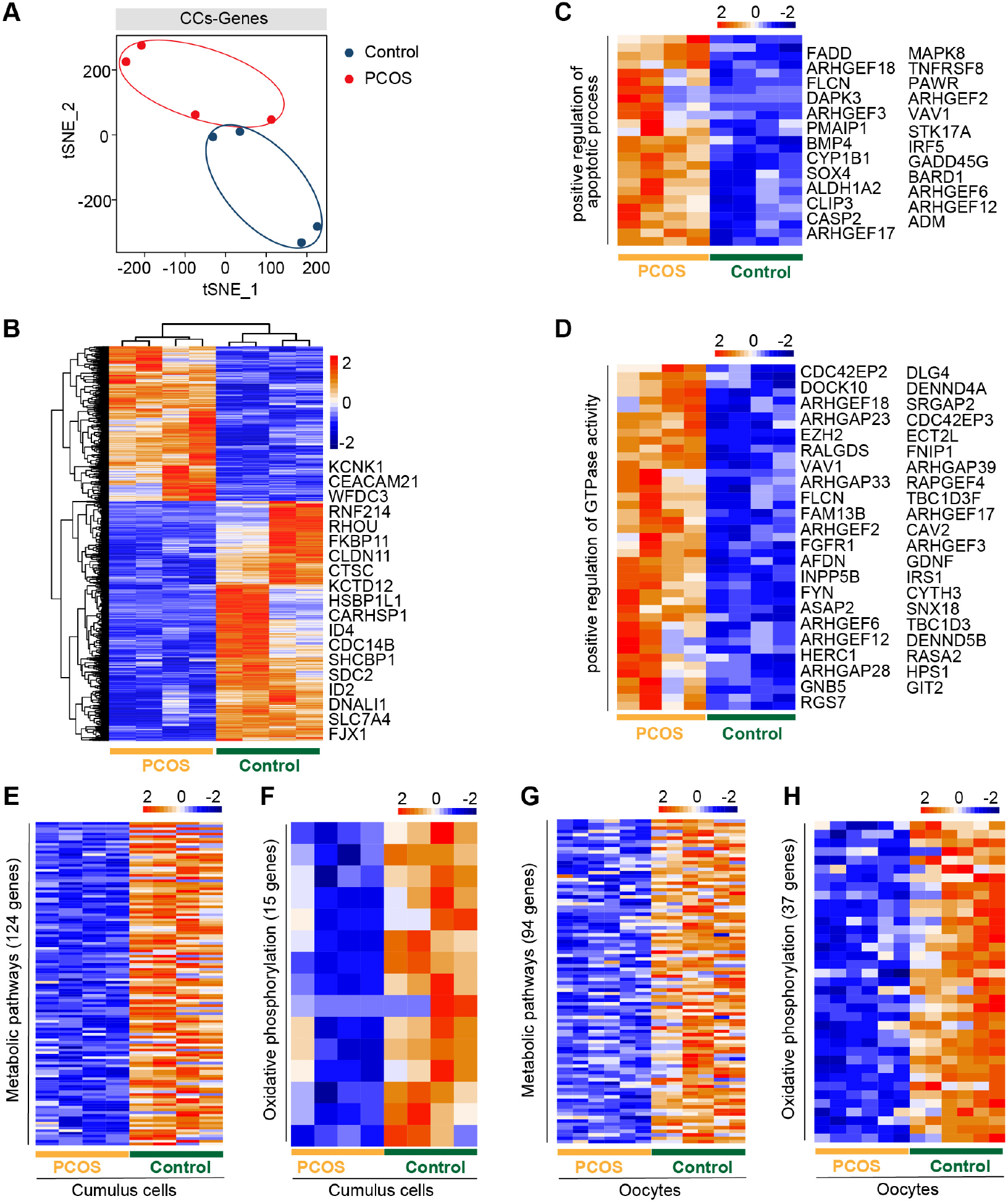
Gene expression pattern differs between PCOS and Control CCs. (**A**) Visualization of gene expression by RNA-seq from four CC samples by t-SNE, showing that CCs are clustered into two distinct subpopulations. Red and blue points represent PCOS and Control CCs, respectively. (**B**) Heatmap of differentially expressed genes (DEGs) in CCs between PCOS and Control. The number of up-DEGs is 704 and down-DEGs is 1,091. Color key from blue to red indicates relative gene expression levels from low to high, respectively. (**C**) Heatmap of genes involved in positive regulation of apoptotic process upregulated in PCOS CCs. (**D**) Heatmap of genes related to positive regulation of GTPase activity upregulated in PCOS CCs. (**E**) Heatmap of genes involved in metabolic pathways downregulated in PCOS CCs. (**F**) Heatmap of genes involved in oxidative phosphorylation downregulated in PCOS CCs. (**G**) Heatmap of genes related to metabolic pathways downregulated in PCOS oocytes. (**H**) Heatmap of genes related to oxidative phosphorylation downregulated in PCOS oocytes.

Moreover, downregulated DEGs included oxidative reduction and glycogen biosynthetic processes, and response to hypoxia (Supplementary Fig. S2B). Decreased expression levels of “response to estrogen” related genes may explain patients with PCOS less sensitive to estrogen and increased T levels (Supplementary Table SI), supporting the increased androgen biosynthesis in PCOS theca cells and inhibition of aromatase in granulosa cells ^29,30^.

By KEGG analysis of crucial signaling pathways, strikingly, among these signaling pathways (Supplementary Fig. S2C, D), many genes involved in metabolic and oxidative phosphorylation pathways showed decreased expression in CCs from women with PCOS (Fig. 2E, F), consistent with those of oocytes from PCOS women (Fig. 2G, H). However, the genes of metabolic pathways and oxidative phosphorylation were not the same in CCs and oocytes owing to their different cell types. Some genes involved in PI3K-Akt signaling pathway displayed high expression levels in CCs from PCOS patients (Supplementary Fig. S2E). In addition, genes related to MAPK and Ras signaling pathway also were expressed at increased levels in PCOS CCs (Supplementary Fig. S2F, G). Our analysis revealed simultaneous dysfunction of oxidative phosphorylation and metabolic pathways in both CCs and oocytes, in addition to the changes in previously known signaling pathways found in granulosa cells or cumulus cells.

### Disorder of mitochondrial function and communication in PCOS oocytes and CCs

Bidirectional communication between oocytes and their associated somatic cells plays a pivotal role in fertility and embryogenesis ^31^. To investigate whether the crosstalk is dysfunctional between oocytes and CCs in PCOS, we explored signaling pathway that are potentially involved in the interaction between oocytes and CCs. NOTCH signaling pathway, such as ligands DLL3 and JAG2 had no significant difference in oocytes between PCOS and Controls, and only receptor NOTCH3 and target gene HES1 were differentially expressed in PCOS CCs (Supplementary Fig. S3A). In terms of the gap junction, expression of three key genes related to this pathway including GJC1, GJA3 and GJA1, did not differ in oocytes/ CCs between PCOS and Controls (Supplementary Fig. S3B). The ligand KITLG and the receptor KIT displayed increased expression levels in CCs and oocytes from PCOS patients, respectively (Supplementary Fig. S3C).

Remarkably, the ligand GDF9 in oocytes was downregulated in the TGF-β signaling pathway, whereas the ligand BMP4 was upregulated in CCs of PCOS patients. Although expression of receptors did not differ in CCs, target genes correlating with cumulus–oophorus extracellular matrix and luteal function, including TNFAIP6, PTX3, ID1, ID2, ID4, were downregulated in PCOS CCs (Fig. 3A). This suggests incomplete cumulus expansion and aberrant luteinization, which presumably affects ovulation in women with PCOS. Overall, the bidirectional communication between the oocytes and CCs may be disrupted.

**Figure 3.**
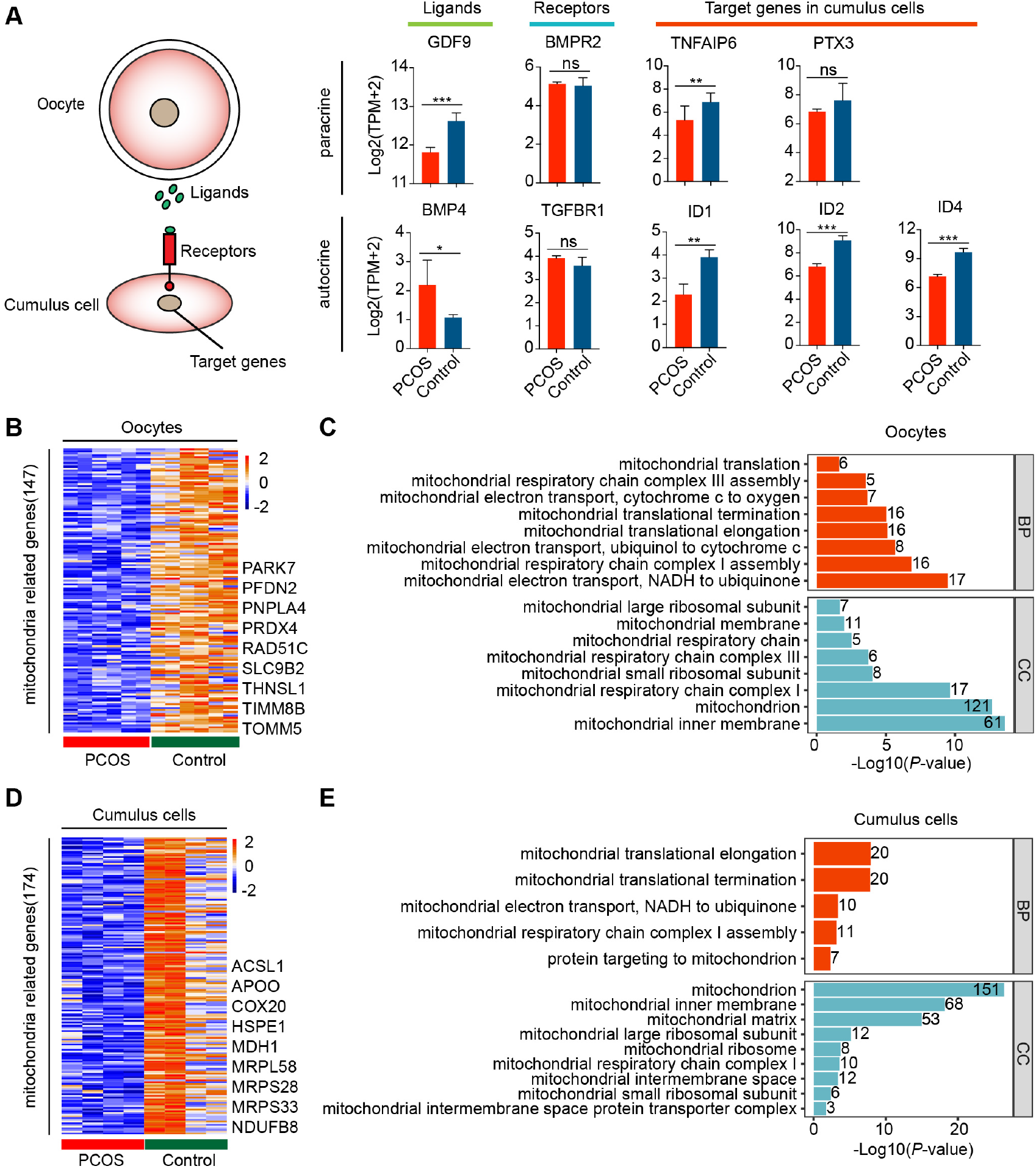
Disorder of mitochondrial function and communication in CCs and oocytes with PCOS. (**A**) TGF-β signaling pathway is involved in the Oocyte-GC interactions. The schematic diagram at the left shows the relationship among these genes. Diagrams at the right show the relative expression levels (log_2_ [TPM+2]) of ligands, receptors, and target genes between PCOS and Control in oocytes (n = 3, participants) and CCs (n = 2, participants). Data represents mean ± SD. *p < 0.05, **p < 0.01, ***p < 0.001, ns, not significant. (**B**) Heatmap of differentially expressed genes involved in mitochondria that were downregulated in oocytes with PCOS. (**C**) GO enrichment analysis of downregulated mitochondria genes in PCOS oocytes. Red bars represent Biological Process and blue bars represent Cellular Component. (**D**) Heatmap of differentially expressed genes involved in mitochondria downregulated in PCOS CCs. (**E**) GO enrichment analysis of downregulated mitochondria genes in PCOS CCs.

Above functional enrichment analysis of oocytes and CCs demonstrated alterations in genes related to metabolism and oxidative phosphorylation. Furthermore, 147 of genes involved in mitochondrial function were downregulated in PCOS oocytes (Fig. 3B, Supplementary Table SVII). Also, 174 genes associated with mitochondrial function were downregulated in PCOS CCs (Fig. 3D; Supplementary Table SVIII), and CCs and oocytes shared 18 genes related to mitochondria. Downregulation of mitochondria-related genes such as NDRG4, UCP2, MRPS26 by RNA-seq was further validated by real-time qPCR (Supplementary Fig. S3D). We also investigated mitochondrial function and components by GO analysis, including Biological Process (BP) and Cellular Component (CC). Downregulated mitochondrial genes in oocytes and CCs were enriched for the same CC terms, including mitochondrial inner membrane, mitochondrial large ribosomal subunit, mitochondrial respiratory chain complex I and mitochondrial small ribosomal subunit (Fig. 3C, E). Furthermore, several BP terms also were enriched in oocytes and CCs, including mitochondrial translational elongation /termination, mitochondrial respiratory chain complex I assembly and mitochondrial electron transport, NADH to ubiquinone. Globally, GO analysis of mitochondria related genes highlighted that mitochondrial dysfunction and faulty mitochondrial components simultaneously occur in oocytes and CCs in women with PCOS, which might contribute to oocytes with decreased fertilization and impaired embryonic development.

### Expression pattern of TEs in oocytes between PCOS and Control

In addition to alteration of gene expression in oocytes and CCs, we also investigated whether transposable elements (TEs) are involved in the pathogenesis of PCOS. Two clusters representing PCOS or Control were clearly identified (Fig. 4A), and PerMANOVA analysis also demonstrated that expression pattern of TEs can distinguish PCOS from Control oocytes (Supplementary Table SIV, *P* =0.004). PCOS oocytes showed 455 upregulated and 371 downregulated TEs (Fig. 4B; Supplementary Table SIX). Most upregulated or downregulated TEs in PCOS oocytes belonged to LTR elements, LINEs, and SINEs (Supplementary Fig. S4A). LTR elements accounted for the largest proportion of main classes. Proportion of downregulated TEs (43.08%) classified in LTR elements was slightly higher than upregulated TEs (39.08%). In terms of superfamily, the differentially expressed TEs in PCOS oocytes mainly pertained to ERV1, ERVL-MaLR, Alu, L1, hAT-Charlie. Upregulated TEs belonged to ERV1 and ERVL-MaLR, in contrast to L1, Alu and hAT-Charlie (Supplementary Fig. S4B). We further analyzed LTR retrotransposons which are proportionately greater than other repeat classes. Approximately 24% of ERV1 elements were highly expressed in PCOS oocytes, including LTR1C, LTR16C, MLT1J2, MER4C, MLT1G1, in contrast to the 11% downregulated expression of ERV1 elements, including LTR12C, MLT1H1, MER41B, LTR7Y (Supplementary Fig. S4C).

**Figure 4.**
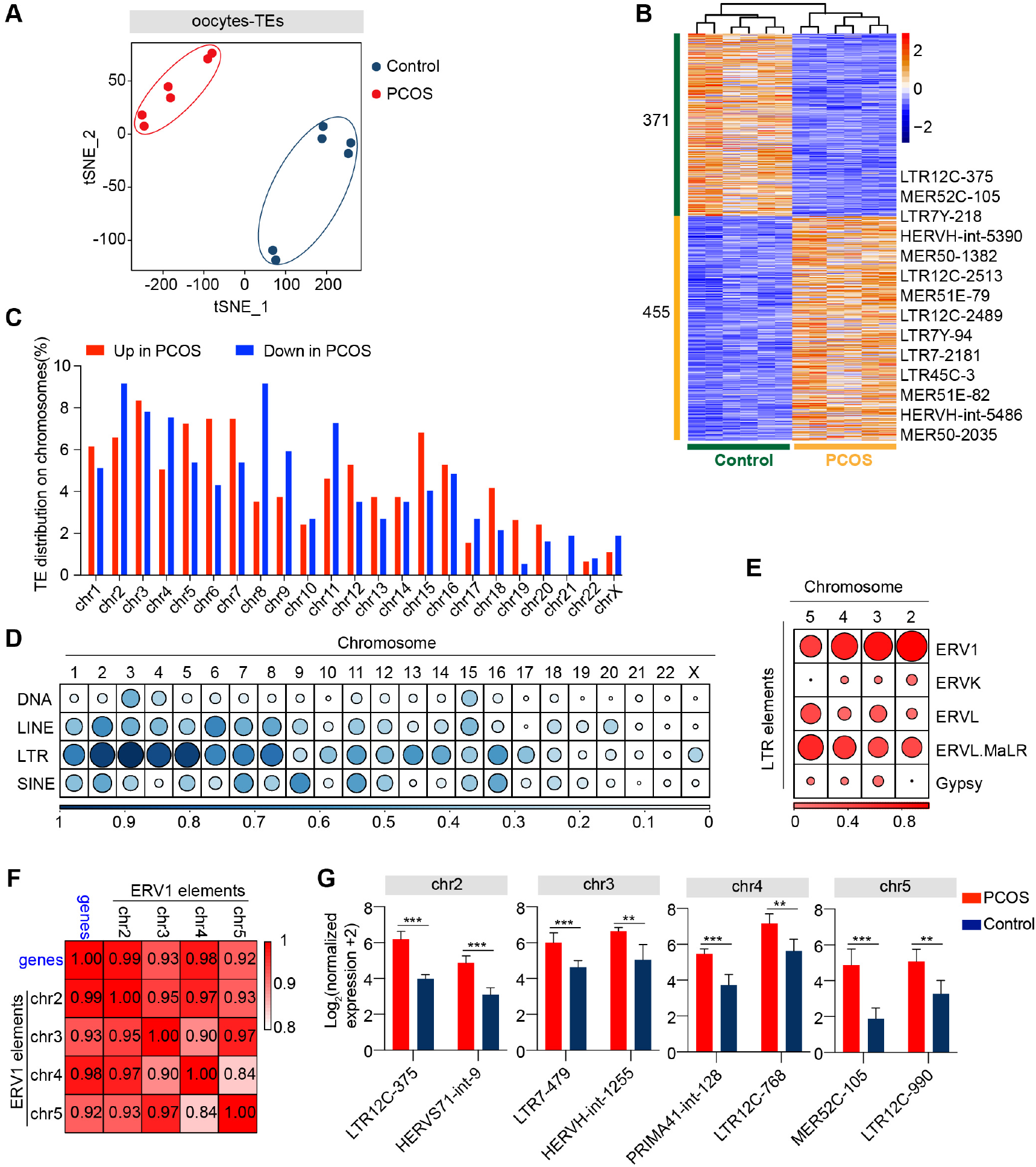
Retrotransposons in PCOS and Control oocytes. (**A**) Visualization of TE expression of oocytes by t-SNE, showing two clusters, including one for PCOS-oocytes and the other for Control-oocytes (n=6). Each dot corresponds to one oocyte, clustered by color. (**B**) Heatmap of differentially expressed TEs (DETs) in oocytes between PCOS and Control. Number of up-DETs is 455 and down-DETs is 371. Color key from blue to red indicates relative gene expression levels from low to high, respectively. (**C**) Proportion of differentially expressed TEs in each chromosome. Red bars represent percentage of up-regulated TEs and blue bars represent percentage of down-regulated TEs. (**D**) Proportion of DETs classified by repeat classes on different chromosomes. Dots from large to small or color from dark to light represent proportion from high to low. (**E**) Proportion of differentially expressed LTR elements classified by repeat super-families on chromosome 2, 3, 4, 5. (**F**) Correlation between expression levels of ERV1 elements on chromosomes 2, 3, 4, 5 and of protein coding genes in PCOS oocytes. Correlation coefficient is shown on the plot. (**G**) Expression levels of differentially expressed ERV1 elements on chromosome 2, 3, 4, 5 in PCOS oocytes, and shown are two differentially expressed ERV1 elements on each chromosome.

To analyze the chromosomal distribution of differentially expressed TEs, we classified differentially expressed TEs according to the chromosome position (chromosome 1-22, X chromosome) (Fig. 4C). Differentially expressed TEs were not uniformly dispersed in the human chromosomes. Most differentially expressed TEs were distributed on chromosomes 1 to 8 chromosome. In contrast, TEs on chromosomes 20, 21, 22, and X chromosomes were less differentially regulated (Supplementary Fig. S4D). Moreover, differentially expressed TEs were mainly enriched in LTR elements on chromosomes 2, 3, 4, and 5 (Fig. 4D). The LTR elements were composed of Gypsy, Copia, Bel-Pao, Retrovirus and ERV, based on the classification system for TEs ^12^. Notably, most differentially expressed LTR elements on chromosome 2, 3, 4, and 5 were classified as ERV1 elements (Fig. 4E).

To confirm whether these differentially expressed ERV1 elements were involved in the occurrence of PCOS, we screened 13 most significantly upregulated genes with - log_10_ (padj) > 10 from the differentially expressed genes between PCOS oocytes and Controls (Supplementary Table SX), and then performed correlation analysis by Pearson’s between the average normalized expression levels of 13 most significantly PCOS-specific protein-coding genes and of the upregulated ERV1 elements on chromosome 2, 3, 4, 5 in PCOS oocytes. Strikingly, highly upregulated ERV1 elements on chromosome 2, 3, 4, 5 are significantly correlated with those of 13 protein-coding genes notably including tubulin associated genes *TUBA1C*, *TUBB8P8* and *TUBB8* (Fig. 4F) and these ERV1 elements were activated in PCOS oocytes (Fig. 4G), suggesting that ERV1 elements may be involved in the alterations in gene regulation observed in PCOS.

### TE expression profile in CCs between PCOS and Control

We wondered whether TEs also are differentially expressed in PCOS CCs. By t-SNE clusters, transcription of TEs in PCOS CCs clearly was distinguishable from that of Control (Fig. 5A). Overall alteration of TE expression profile in PCOS CCs was not as dramatic as in PCOS oocytes and TE expression profile in PCOS CCs did not differ from that of Controls by PerMANOVA analysis (Supplementary Table SIV, *P* =0.280). Compared with Control CCs, 14 differentially expressed TEs displayed upregulation and 7 downregulation in PCOS CCs (Fig. 5B, Supplementary Table SXI). Most differentially expressed TEs were enriched in L1 elements (Supplementary Fig. S4E and Fig. 5C). L1 retrotransposons are autonomously active and can integrate into the genome, which can remodel gene structure and further impact human evolution and disease ^32^. Most differentially expressed L1 elements exhibited high expression levels in PCOS CCs, containing L1MC1, L1MB3, L1MEf, L1ME1 and L1MC4 (Fig. 5D). The role CC TEs play in PCOS remains to be determined, though we cannot rule out the possibility that TEs may influence transcription of genes implicated in PCOS.

**Figure 5.**
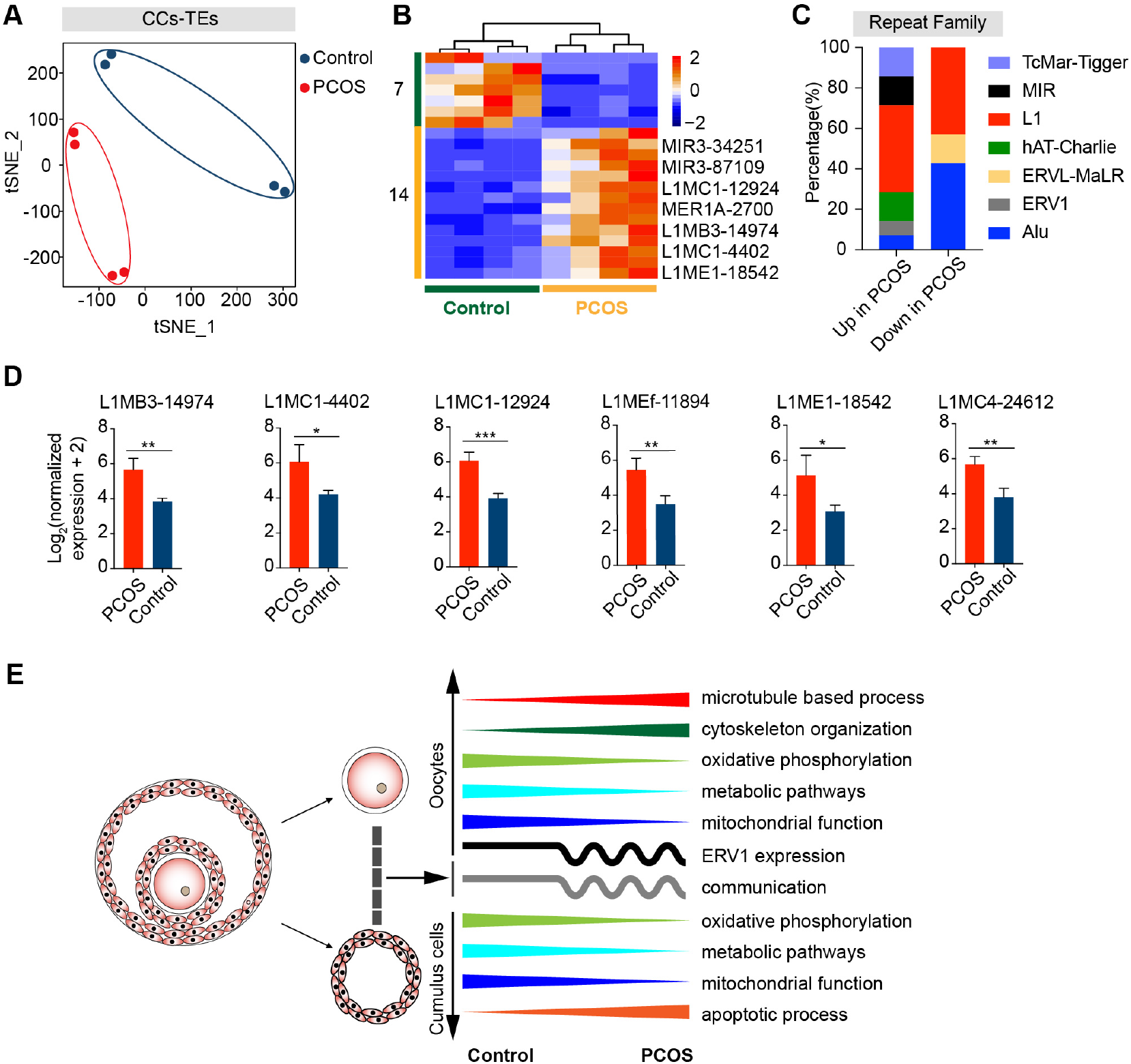
Retrotransposons in PCOS and Control CCs. (**A**) Visualization of TE expression of CCs by t-SNE, showing two clusters, including one for PCOS-CCs (four samples) and the other for Control-CCs (four samples). Each dot corresponds to CCs. (**B**) Heatmap of all differentially expressed TEs (DETs) in CCs between PCOS and Control. Number of up-DETs is 14 and down-DETs is 7. (**C**) Percentage of upregulated or downregulated TEs classified by repeat superfamily. (**D**) Expression levels of differentially expressed L1 elements in PCOS CCs. (**E**) Summary of molecular features in oocytes and CCs in the occurrence of PCOS.

## Discussion

By simultaneous transcriptome analysis of the oocytes and CCs from the same PCOS patients, compared with controls at similar age, we identified differences in global gene expression patterns and in expression of TEs in oocytes or CCs in PCOS patients. Our data revealed surprising findings of altered microtubule-related genes, in addition to genes including metabolisms, as reported previously. We also discovered changes in the transcription of retrotransposons, which would be likely to influence the transcriptome of PCOS oocytes. ERV1 elements particularly may be involved in the pathogenesis of PCOS. In contrast, CCs from PCOS patients contained fewer alterations in TEs. This data also supports the notion that most retrotransposon events occur in parental germ cells and only rarely in somatic cells ^33,34^.

Previous studies on the transcriptomic profiles of oocytes and granulosa cells or CCs from PCOS patients have shown molecular abnormalities present in PCOS patients. Wood *et al.* reported that several differentially expressed genes of PCOS oocytes are related to spindle dynamics and centrosome function ^6^. A similar study using single-cell RNA-seq found that some genes involved in meiosis and gap junction exhibited abnormal expression in PCOS oocytes ^7^. Here, we find that genes involved in microtubule-based processes, cytoskeleton organization were differentially expressed in PCOS oocytes. Specifically, crucial genes involved in microtubule based processes, *TUBB8* and *TUBA1C,* were overexpressed in PCOS oocytes. These changes would be expected to negatively affect oocyte maturation and subsequent embryonic development. Interestingly, mutations in *TUBB8* contributed to the disruption of oocyte meiotic spindle assembly and maturation and led to female infertility ^21^. Although PCOS patients have more oocyte yield from stimulation during IVF cycle, poor oocyte quality causes lower fertilization and implantation rates, poor quality of embryos, decreased pregnancy and increased miscarriage rates ^35–43^. Indeed, oocytes from women with PCOS displayed maturation arrest and disrupted spindles after IVM as shown in our study. Hence, increased expression of *TUBB8* and *TUBA1C* resulting in defective microtubule and spindle formation could be one major factor contributing to a decline in oocyte quality in PCOS women.

In addition, crucial signaling pathways are altered in oocytes from PCOS women. For instance, genes related to MAPK signaling pathway show decreased expression in oocytes from women with PCOS, and MAPK plays a pivotal role in regulating oocytes meiotic resumption ^44,45^. Expression of genes involved in mTOR signaling also is decreased in oocytes from PCOS women. The mTOR–eIF4F pathway spatiotemporally regulates chromosome segregation and functional spindle formation during meiosis in mammalian oocytes ^25^. However, it is unclear whether aberrant MAPK and mTOR are related to the overexpression of microtubule-associated genes in PCOS.

Our findings also imply potential roles of TEs in PCOS oocytes and CCs. The pattern of TE expression distinguishes PCOS from control oocytes. Most differentially expressed TEs are classified as LTR elements, belonging to the superfamilies ERV1, ERVL-MaLR, Alu, L1, hAT-Charlie. Also, ERV1 elements differentially expressed on chromosome 2, 3, 4, 5 in oocytes may be associated with PCOS. Future studies should determine whether this serves as a biomarker of PCOS. In addition, fewer TEs are altered in CCs than in oocytes. It will be interesting to determine whether these L1s contribute to genome instability in women with PCOS.

Most of previous studies on PCOS focused on ovarian somatic cells, peripheral blood or other cell types. For instance, cumulus cells of PCOS patients display abnormal characteristics of gene expression including dysregulated growth factors, steroid metabolism, cell cycle, steroid hormone biosynthesis and hypomethylated genes related to the synthesis of lipid and steroid ^7–9,46^. Dysregulation of inflammatory function also has been found in PCOS patients through transcriptome and DNA methylation analysis of peripheral blood and granulosa-lutein cells ^47–49^. In addition, aberrant expression of MicroRNAs in granulosa cells, theca cells and follicular fluid might be involved in the development of PCOS ^50–52^. In our study, global gene expression in CCs from PCOS women with also differs from controls. Furthermore, two key genes are differentially expressed in CCs from PCOS patients including LH/choriogonadotropin receptor (LHCGR) and insulin receptor gene (INSR) (Supplementary Table 6), which were identified as susceptibility genes for PCOS in previous GWAS and strongly associated with anovulation ^4,5,53^. Functional annotation of genes upregulated in PCOS CCs shows alteration in the positive regulation of GTPase activity, apoptotic process and the steroid metabolic process. Alteration of gene expression related to ‘positive regulation of apoptotic process’ may activate apoptosis of CCs and indirectly reduce the quality and developmental competence of the oocytes ^54^. Conversely, genes related to the carbohydrate metabolic process and response to lipopolysaccharide show decreased expression. Abnormal metabolism in CCs is thought to contribute to the clinical features shared among PCOS, obesity and diabetes. In support, PCOS women have increased prevalence of metabolic syndrome, impaired glucose tolerance and obesity ^55–58^. Also, ovarian reserve may be affected by obesity in women variably depending on presence of PCOS ^59^. Meanwhile, some signaling pathways were also dysregulated in CCs from PCOS patients, such as PI3K-Akt and MAPK signaling pathway, which are known related to PCOS, involving insulin resistance and excessive androgen production ^60–63^.

Our finding reveals that metabolic pathways and oxidative phosphorylation are dysregulated in both oocytes and CCs of PCOS patients, further validating and explaining the phenotype that PCOS women exhibit increased risk of metabolic syndrome. Anovulation is a common cause of infertility in PCOS patients and some studies have shown that anovulatory infertility is related to metabolic abnormalities ^64–66^. Thus, for women with anovulatory PCOS, abnormal metabolism in CCs may be related to anovulation. Moreover, disordered mitochondrial function in oocytes and CCs may contribute to declined oocyte developmental competence. Mitochondria are essential for oocyte development potential and oocyte rejuvenation ^67^, and mitochondrial dysfunction in oocytes is found in women with PCOS ^68,69^. Mitochondrial functions may be prematurely activated at GV stage oocytes of PCOS ^70^, and the oocytes may have impaired mitochondrial ultrastructure and functions, including compromised inner mitochondrial membrane potential and electron transport chain ^71^, consistent with our results. Crosstalk between oocytes and CCs also can be perturbed by alteration of the ovarian microenvironment, including oxidative stress caused by mitochondrial respiratory dysfunction, which leads to enhanced ROS production ^72^. Additionally, our analysis reveals that communication between oocytes and CCs may be disrupted in PCOS patients, notably members of the TGF-β superfamily, including BMPs/GDFs that are important regulators in human folliculogenesis and ovulation ^73^. It is likely that the abnormalities of essential signaling pathways may attribute to the disorder of CCs and further influence the quality of oocytes in PCOS patients.

## Conclusion

Taken together, our comprehensive analysis of oocytes and CCs, including global gene expression and TEs expression profiling, reveals unique molecular features of PCOS (Fig. 5E). Furthermore, new candidate genes and TEs in oocytes and CCs could serve as the signatures of PCOS. Increased expression levels of TUBB8 and TUBA1C and resultant spindle defects can specifically define oocyte quality of PCOS patients. Overall, these findings may suggest future treatment strategies to improve oocyte maturation and developmental competence in PCOS women.

## Materials and methods

### Ethical approval

Informed consents were obtained from all participants included in the study. This study was approved by the Ethics Committee of TianJin Medical University General Hospital (No: IRB2018-102-01) and conducted in accordance with approved institutional guidelines.

### Human subjects

The study subjects included five women without PCOS (Controls) with BMI between 17.70 and 23.50 kg/m^2^ and five PCOS patients with BMI between 19.00 and 28.10 kg/m (*P* = 0.114). The average age of PCOS patients and Controls were 32.40 ± 1.29 and 35.60 ± 2.23, respectively (*P* = 0.249). The demographic and clinical characteristics of all participants including age, BMI, LH, and others, were collected and summarized in Supplementary Table SI (available online).

PCOS patients were diagnosed according to the Rotterdam criteria ^74,75^, which meets two of the following three features, Oligo- or anovulation, clinical and/or biochemical signs of hyperandrogenism, and polycystic ovary by ultrasound. We excluded other etiologies, such as congenital adrenal hyperplasia, androgen-secreting tumors, and Cushing’s syndrome. Women included in the Control group had regular menstrual cycles, normal sonographic appearance of ovaries, and no diabetes or clinical signs of PCOS.

### Ovarian stimulation and oocyte retrieval

All participants underwent controlled ovarian stimulation using the GnRH antagonist protocols with the recombinant human FSH (rhFSH) and ICSI for male fertility. An ultrasound scan and serum estradiol assays were performed for monitoring follicular size. When two or more follicles were at least 12 mm in diameter, 10,000 IU human chorionic gonadotropin (hCG) was administered 36 hours before oocyte retrieval. The amount and duration of rhFSH treatment were similar in both PCOS patients and Controls, exhibiting no statistically significant difference (Supplementary Table SI).

### Isolation of single oocyte and CCs

Cumulus-oocyte complex (COC) was isolated via ultrasound-guided vaginal puncture and classified according to the oocyte nuclear maturation stage: GV (germinal vesicle), MI (metaphase I) and MII (metaphase II). We only collected GV stage oocytes and surrounded CCs for this study, whereas MII stage oocytes were used for clinical fertilization.

The CCs were collected as previously described ^8^. Briefly, CCs were mechanically stripped from oocytes shortly under stereomicroscopy prior to ICSI and then isolated CCs were dispersed into single cell with 0.03% hyaluronidase (H6254-500MG, Sigma-Aldrich) and resuspended three times using PBS. Separated CCs were counted up to 500 cells and placed in lysate. Tyrode’s Acidic Solution (T1788-100ML, Sigma-Aldrich) was used to facilitate stripping the zona pellucida to produce naked oocytes. Oocytes were observed under a microscope to ensure absence of contamination with CCs. Naked oocytes were carefully washed three times using PBS with 0.1% polyvinylpyrrolidone (PVP, P0930-50G, Sigma-Aldrich) to prevent sticking to handling tools or dishes, and then placed in lysate.

Six oocytes and four CC samples were collected from five PCOS patients and six oocytes and four CC samples from five Controls, respectively. Oocytes and CCs from three PCOS patients and three Controls were used for RNA-seq, and CCs for PCOS-3 patient and Control-3 were lost during collection. Oocytes from two PCOS patients and two Controls were used for subsequent immunofluorescence microscopy or *in vitro* maturation (Supplementary Table SII).

### Library construction from oocytes or CCs and sequencing

Individual oocytes or 500 CCs were transferred into lysis buffer quickly and Smart-seq2 protocol ^76^ was used to synthesize cDNA for single cell RNA-seq analysis. After reverse transcription of mRNA and amplification of cDNA, qPCR analysis was performed to check the quality of the cDNA libraries using housekeeping gene, GAPDH. Variation in the expression of GAPDH was minimal, and negative control (NTC) did not detect any product (Supplementary Fig. S1A). The RNA-Seq libraries were constructed by TruePrep DNA Library Prep Kit V2 for Illumina^®^ (TD503-02, Vazyme Bio-tech) following the instruction manual. Meanwhile, to ensure the accuracy and repeatability of RNA-seq data, we performed a duplicate when we constructed library for every oocyte, and collected CC sample to match with the retrieval oocytes from two PCOS patients and two Controls, so each donor had two CC samples for RNA-seq. We performed 14 cycles of PCR to amplify the cDNA library and simultaneously barcoded it. Final indexed libraries were pooled and sequenced on an Illumina HiSeq X10 platform with a 150-bp paired-end read length.

### RNA-Seq data processing and analysis

RNA-seq raw reads with low-quality bases, adapters were trimmed by Trimmomatic ^77^ to obtain clean reads (parameters: -PE-phred33-SLIDINGWINDOW:4:15 - LEADING:10-TRAILING:3-MINLEN:36). Clean reads were aligned to the UCSC human hg19 reference using the Hisat2 with default settings ^78^. Read counts of each gene annotated in RefGene were calculated by featureCounts with default parameters ^79^. RNA-seq libraries of each oocyte and each CC sample were sequenced at an average depth of approximately 4.6 million reads per oocyte and 6.8 million per CC sample (Supplementary Fig. S1B) and the average ratio mapped for each oocyte and each CC sample was 72.98% and 54.88%, respectively (Supplementary Fig. S1C). The read counts were loaded into RStudio (v3.6.1). For the accuracy of gene expression levels, only genes with TPM > 1 in at least one oocyte or CC sample were analyzed. DESeq2 ^80^ was used to obtain statistical significance of differentially expressed genes between PCOS and Control. Only the genes with a fold change of log_2_ transformed larger than log_2_(1.5) and adjusted *P* value < 0.05 from DEseq2 results were considered to be differentially expressed. Adjusted *P* values were computed in DESeq2 using the Wald test, adjusted for multiple testing using the procedure of Benjamini and Hochberg ^81^. Gene ontology (GO) and KEGG analysis of differentially expressed genes was performed using DAVID (v6.8) ^82^, and only enriched pathways that showed *P* value < 0.05 were chosen as significant enrichment. Gene expression level in a sample was quantified as the transcripts per million (TPM), which was calculated according to the following formula: 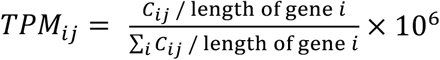, where *C_ij_* was the count value of gene *i* in sample *j*. Genes with expression transformed to the TPM values were used for t-SNE dimension reduction by R package ‘Rtsne’ and t-SNE map drawn using the R package ‘ggplot2’. Diagrams were generated by TPM value of each gene that were plus two then log_2_ transformed.

### Transposable elements analysis

Clean reads were aligned to the UCSC human hg19 reference by STAR with parameters ‘-winAnchorMultimapNmax 100’ and ‘-outFilterMultimapNmax 100’ ^83^. Based on the previous method ^84^, only the TEs mapping their distributions in intergenic regions were considered, excluding the location between the transcription start sites and transcription end sites of genes. TEs annotated in UCSC Genome Browser (RepeatMasker) were counted using featureCounts. The reads mapped to

TEs of each oocyte and each CC sample were sequenced at an average depth of approximately 0.7 million reads per oocyte and 2.0 million per CC sample (Supplementary Fig. S1D) and the average ratio mapped of each oocyte and each CC sample were 11.55% and 16.58%, respectively (Supplementary Fig. S1E). The DESeq2 was subsequently applied to identify differentially expressed transposable elements, and only the transposable elements with a fold change of log_2_ transformed larger than log_2_(1.5) and adjusted P value < 0.05 from DEseq2 results were considered to be differentially expressed, and expression of TEs were normalized by DESeq2. The normalized expression of TEs was used as input for t-SNE dimension reduction.

### Correlation analysis

We screened genes with −log_10_ (padj) > 10 from the differential gene expression of oocytes, showing the most significantly upregulated protein-coding genes in PCOS oocytes, and then calculated the average normalized expression of these genes in each oocyte (*G_i_*). By counting the proportion of differentially expressed TEs on each chromosome according to the classification of the classes and super-families, we found out the super-families significantly enriched and upregulated in PCOS oocytes, and then calculated the average normalized expression of the super-families in each oocyte *(T_i_*). Log_2_ transformed *G_i_* values and log_2_ transformed *T_i_* values were used as inputs for correlation analysis using Pearson’s correlation. The plot was drawn by the R package ‘pheatmap’.

### Oocyte in vitro maturation (IVM)

*In vitro* maturation of germinal vesicle oocytes was achieved using SAGE IVM media kit (ART-1600). Briefly, germinal vesicle oocytes surrounded by CCs, retrieved during IVF cycles, were washed twice with SAGE washing medium and then cultured in SAGE IVM medium with 75 IU HMG (human menopausal gonadotropin) for approximately 24 h under paraffin oil at 37°C in high humidified atmosphere of 6% CO2 in air ^85^. Oocyte maturation was assessed by the presence of the first polar body using the inverted microscope. For immunofluorescence microscopy of oocytes, surrounding CCs were removed by hyaluronic acid.

### Immunofluorescence and confocal microscopy

Immunofluorescence microscopy of oocytes for spindle imaging was performed based on the previous method ^86^. Oocytes were fixed in fixative (MTSB XF) at 37 °C for at least 30 min and then washed four times with washing buffer (phosphate-buffered saline, supplemented with 0.02% NaN3, 0.01% Triton X-100, 0.2% non-fat dry milk, 2% goat serum, 2% bovine serum albumin and 0.1 M glycine). Afterwards, oocytes were left in washing buffer for 2 hr at 37 °C for blocking. To determine protein expression, oocytes were incubated with anti-TUBB8 antibody (1:500, SAB2700070,

Sigma-Aldrich) or anti-TUBA1C (1:300, PA516891, Thermo Scientific) overnight at 4 °C. Oocytes were washed and incubated with secondary goat anti-rabbit IgG Alexa Fluor 594 antibody (1:200, 111-585-003, Jackson) at 37 °C for 2 h and the Hoechst 33342 (1:200, H3570, Life Technologies) to label DNA. Oocytes were mounted on glass slides and sealed with nail polish, and examined with a confocal laser-scanning microscope (Leica). Oocytes not incubated with anti-TUBB8 or TUBA1C antibodies but only with the secondary antibody and stained DNA with Hoechst 33342 served as control for non-specific staining. The control oocytes showed only DNA staining (in blue) and no other non-specific staining.

### Gene expression analysis by Real-Time quantitative PCR

mRNA of CCs was reverse transcribed to cDNA according to the Smart-seq2, and the products diluted at the final concentration of 0.25ng/ μl. Real-time quantitative PCR (qPCR) reactions were performed in duplicate with the FS Universal SYBR Green Master (4913914001, Roche) and run on the iCycler MyiQ2 Detection System (Bio-Rad). Each sample was repeated three times and analyzed using GAPDH as the internal control. The amplification program was set up as follows, primary denaturation at 95 °C for 10 min, then 40 cycles of denaturation at 95 °C for 15 s, annealing and elongation at 58 °C for 1min, and last cycle for dissociation curve under 55-95 °C. Primers used for qPCR were listed in (Supplementary Table SIII) and confirmed their specificity with dissociation curve.

### PerMANOVA analysis

Differences of global gene or TEs expression between PCOS and Control group were tested by ‘Permutational Multivariate Analysis of Variance’ (PerMANOVA) as implemented by the function adonis in the R package vegan (v2.5-6) based on the Bray-Curtis distance measure (permutation:999).

### Statistical analysis

Data for gene expression levels was analyzed by student’s t-test (paired comparison) and ANOVA (multiple comparisons) using StatView software from SAS Institute Inc. (Cary, NC). Results were represented as mean ± SD, and *P* values for these statistical analyses were based on three oocytes in duplicate or two CCs samples in duplicate from three PCOS patients or three Controls. Data for clinical characteristics and hormone levels between PCOS patients and Controls was analyzed by unpaired student’s t-test using the SPSS 26.0 software and shown as the mean ± SEM (n = 5). Significant differences were defined as \**P*< 0.05, \*\**P*< 0.01 or \*\*\**P*< 0.001.

## Data availability

The accession number for the raw and processed RNA-seq data reported in this paper is GEO: GSE155489

## Authors contributions

J.L. executed experiments, analyzed data, wrote original draft. H.C. was involved in experimental execution and sample collection. M.G., C.T., and H.W. provided critical inputs on data analysis and experimental execution. D.L.K. reviewed the manuscript. X.B. and X.S provided clinical diagnostic information, selected and recruited donors with/without PCOS. L.L. conceptualized the study and reviewed the manuscript.

## Supplementary data

### Supplementary Tables

**Supplementary Table SI: Clinical characteristics and hormone levels of the participants.**

**Supplementary Table SII: Samples for experiments.**

**Supplementary Table SIII: Primers for real-time qPCR.**

**Supplementary Table SIV: PerMANOVA analysis by Bray Curtis distance measure.**

**Supplementary Table SV: Differentially expressed genes of oocytes between PCOS and Control.** Differentially expressed genes of oocytes between PCOS (N=6) and Control (N=6) are included in separate columns.

**Supplementary Table SVI: Differentially expressed genes of CCs between PCOS and Control.** Differentially expressed genes of CCs between PCOS (N=4) and Control (N=4) are included in separate columns.

**Supplementary Table SVII: Mitochondria-related genes differentially expressed in oocytes between PCOS and Control.**

**Supplementary Table SVIII: Mitochondria-related genes differentially expressed in CCs between PCOS and Control.**

**Supplementary Table SIX: Differentially expressed transposable elements in oocytes between PCOS** (N=6) **and Control** (N=6).

**Supplementary Table SX: 13 most significantly upregulated genes out of the differentially expressed genes between PCOS and Control oocytes.**

**Supplementary Table SXI: Differentially expressed transposable elements of CCs between PCOS** (N=4) **and Control** (N=4).

### Supplementary Figures

**Supplementary Figure S1.**
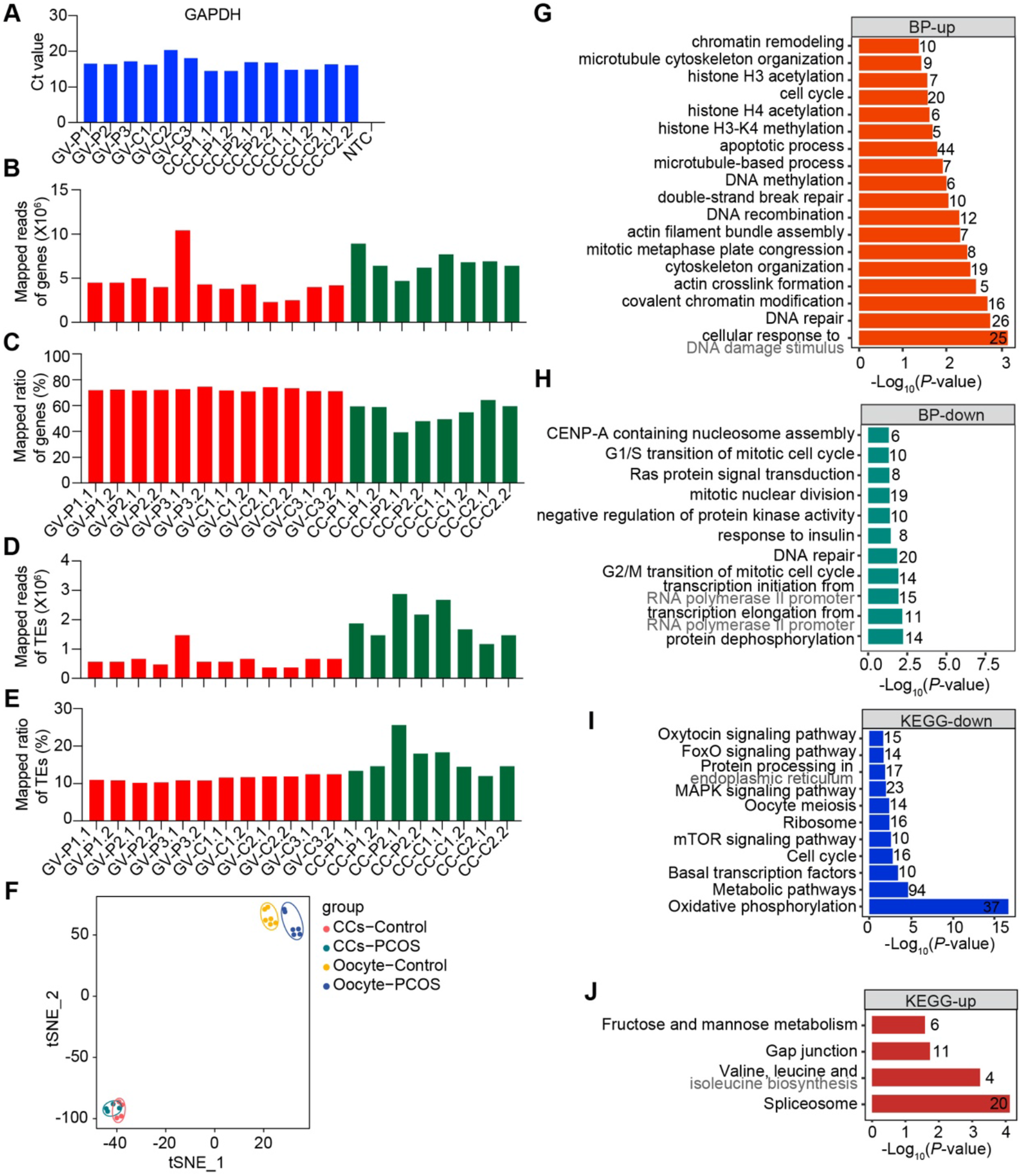
Quality control of RNA-seq analysis and key enrichment analysis of oocytes in patients with PCOS. (**A**) Ct values of GAPDH from oocytes and CCs samples. (**B**) Total mapped reads of genes from oocytes and CCs sample. (**C**) Mapped ratio of genes from oocytes and CCs sample. (**D**) Total mapped reads of TEs from oocytes and CCs sample. (**E**) Mapped ratio of TEs from oocytes and CCs sample. (**F**) Visualization of gene expression of six oocytes and four CCs samples by t-SNE, clustered into four subpopulations including CCs-Control, CCs-PCOS, oocyte-Control and oocyte-PCOS groups. (**G**) Significantly enriched GO terms (biological processes) of DEGs with upregulated expression in PCOS oocytes. (**H**) Significantly enriched GO terms (biological processes) of DEGs with downregulated expression in PCOS oocytes. (**I**) Signaling pathways enriched from DEGs with downregulated expression in PCOS oocytes. (**G**) Signaling pathways enriched from DEGs with upregulated expression in PCOS oocytes.

**Supplementary Figure S2.**
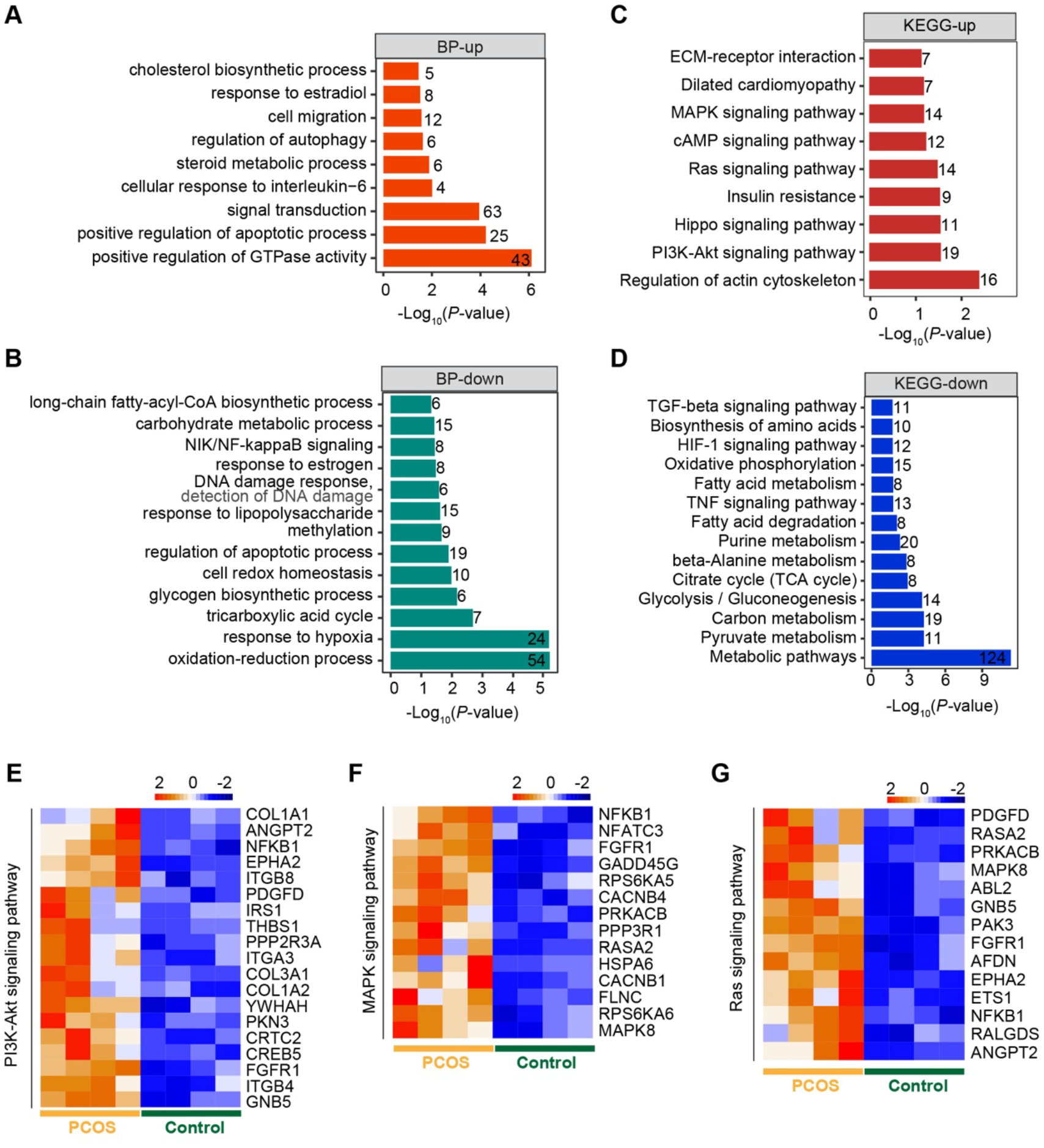
Significantly enriched function and signaling pathways in PCOS CCs. (**A**) Significantly enriched GO terms (biological processes) of DEGs with upregulated expression in PCOS CCs. (**B**) Significantly enriched GO terms (biological processes) of DEGs with downregulated expression. (**C**) Key signaling pathways enriched from DEGs with upregulated expression. (**D**) Key signaling pathways enriched from DEGs with downregulated expression. (**E**) Heatmap of PI3K-AKT signaling pathway upregulated in PCOS CCs. (**F**) Heatmap of genes related to MAPK signaling pathway upregulated. (**G**) Heatmap of Ras signaling pathway upregulated.

**Supplementary Figure S3.**
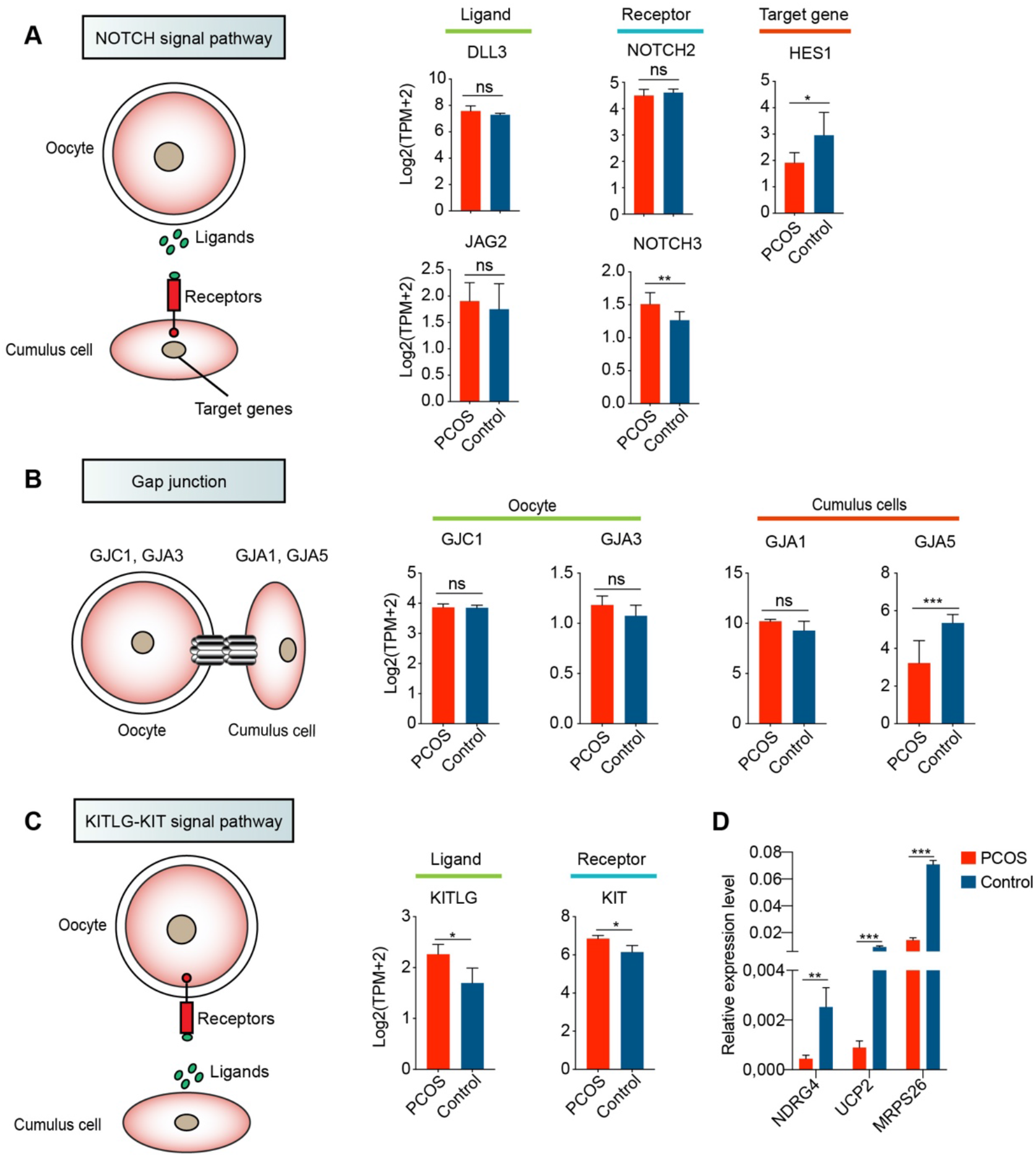
Pathways involved in the Oocyte-CC interactions. (**A**) NOTCH signaling pathway involved in the Oocyte-CC interactions. Schematic diagram on left shows the relationship among these genes. Histograms on right show relative expression levels (log_2_ [TPM+2]) of ligands, receptors, and target genes in oocytes (n = 3, participants) and CCs (n = 2, participants). Red and blue bar represent PCOS and Control, respectively. Data represents mean ± SD. *p < 0.05, **p < 0.01, ***p < 0.001, ns, not significant. (**B**) Gap junction involved in the Oocyte-CC interactions. Histograms on right show relative expression levels of related genes in oocytes and CCs. (**C**) KITLG-KIT signaling pathway involved in the Oocyte-CC interactions. (**D**) Relative expression levels of mitochondria-related genes in CCs from Controls and PCOS patients by qPCR analysis, *p < 0.05, **p < 0.01, ***p < 0.001.

**Supplementary Figure S4.**
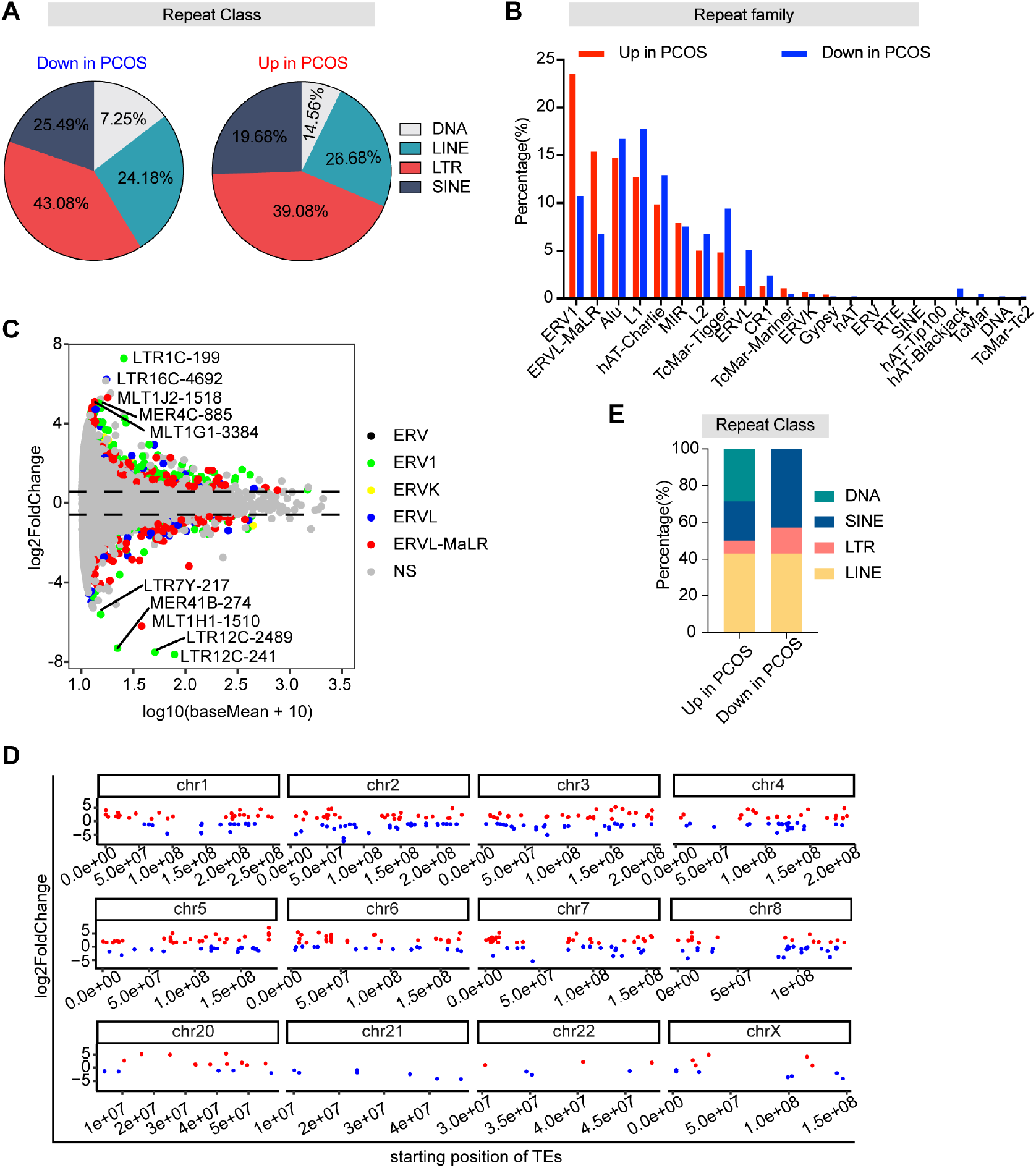
Expression pattern of TEs in oocytes and CCs from PCOS. (**A**) Percentage of all differentially expressed TEs classified by repeat classes in PCOS oocytes. (**B**) Scatter diagram showing differently expressed ERV elements in PCOS oocytes. Different color dots indicate different ERV elements and NS represents the ERV elements with no significant difference. (**C**)Percentage of all differentially expressed TEs classified by repeat family in PCOS oocytes. (**D**) Distribution of differentially expressed TEs in each chromosome. Red and blue dots represent the upregulated and downregulated TEs, respectively. (**E**)Percentage of all differentially expressed TEs classified by repeat classes in PCOS CCs.

